# Sustained learned immunosuppression could not prevent local allergic ear swelling in a rat model of contact hypersensitivity

**DOI:** 10.1101/2025.04.05.647348

**Authors:** Yasmin Salem, Stephan Leisengang, Marie Jakobs, Kirsten Dombrowski, Julia Bihorac, Laura Heiss-Lückemann, Sebastian Wenzlaff, Lisa Trautmann, Tim Hagernacker, Manfred Schedlowski, Martin Hadamitzky

## Abstract

Taste-immune associative learning has been shown to mimic immunopharmacological responses. Conditioned pharmacological effects may therefore be considered as controlled drug dose reduction strategy to maintain treatment efficacy. Against this background, the present study applied an established taste-immune associative learning protocol to a rat model of hapten-induced contact hypersensitivity. After repeated pairings of a saccharin taste (conditioned stimulus, CS) with injections of the immunosuppressant cyclosporine A (CsA, unconditioned stimulus, UCS), animals were sensitized with the hapten. Retrieval started by presenting the CS together with sub-effective doses of CsA. This procedure preserved a conditioned suppression of splenic cytokine production. Compared to full dose treated animals, conditioned effects were neither observed in draining lymph nodes nor did it prevent ear swelling. These findings suggest that local allergic reactions in sensitized animals were most likely too strong to be responsive to the learned immunosuppressive effects. Additionally, symptoms such as itch may be more suited as readout parameter since it better reflects patients’ disease burden. The present study reaffirms that learned immunopharmacological effects can be preserved using a memory-updating approach. It also emphasizes the need to further explore the usability of associative learning protocols in clinical contexts in order to address disease-specific symptoms more effectively.

## Introduction

Allergic contact dermatitis (ACD) is an inflammatory skin disease triggered by daily used objects such as detergents and fragrances that contain reactive low molecular weight chemicals (haptens) (Scheinman et al., 2021). ACD accounts for 30 % of all occupational diseases in industrialized nations (Castanedo-Tardan and Zug, 2012) and it is classified as type IV delayed type allergy. Disease progression involves two distinct phases (Honda et al., 2013). In an initial sensitization phase, hapten exposure leads to a complex formation with self-proteins (haptenization) and a subsequent activation of the innate immune system. Upon re-exposure to the allergen (challenge phase), various activated immune cells and their mediators induce characteristic symptoms such as erythema, oedema and pruritus, caused by the dilation of blood vessels and increased capillary permeability (Dudeck et al., 2011; Honda et al., 2013; Koppes et al., 2017). Specifically, keratinocytes, neutrophils and macrophages secrete cytokines such as tumor necrosis factor (TNF)-α and interleukin (IL)-1), which recruit T cells (Biedermann et al., 2000). These activated T cells, in turn, release other cytokines such as interferon (IFN)-γ, IL-2, IL-17 and IL-5 (Honda et al., 2013; Brites et al., 2020). Additionally, natural killer (NK) T cells produce IL-4, activating B cells and their secretion of immunoglobulin (Ig) M. This process triggers the complement system and subsequently mast cells, which release TNF-α and histamine (Tončić et al., 2011; Brites et al., 2020).

Frequently, the immunosuppressive drug cyclosporine A (CsA) is systemically used to alleviate ACD symptoms by modulating T cell activity (Saary et al., 2005). However, chronic CsA use is accompanied by somatic and neuropsychiatric side effects including nephrotoxicity, depression, and/or anxiety (Mihatsch et al., 1998; Bosche et al., 2015; Brosda et al., 2020), negatively impacting patients’ quality of life (Scheinman et al., 2021). To overcome these drawbacks, learned immunosuppressive placebo responses have been proposed as a strategy to reduce drug dosages while maintaining therapeutic efficacy (Doering and Rief, 2012; Schedlowski et al., 2015). These approaches rely on Pavlovian or classical conditioning and the intense bidirectional communication between the brain and the immune system (Hadamitzky et al., 2020; Hadamitzky and Schedlowski, 2022). Such strategies have been proven effective in experimental animals, healthy subjects and patient populations (Goebel et al., 2002; Wirth et al., 2011; Kirchhof et al., 2018).

In this well-established paradigm, animals are exposed to an unfamiliar gustatory stimulus as conditioned stimulus (CS, saccharin solution) paired with an injection of the immunosuppressive drug CsA as unconditioned stimulus (UCS). Subsequent presentation of the CS alone at a later time point induces both behavioral and immunological changes. These include conditioned taste avoidance (CTA), where animals consume less of the CS, and a suppression of T cell function alongside decreased interleukin (IL)-2 and interferon (IFN)-γ cytokine protein production (Exton et al., 1998; Exton et al., 2002; Pacheco-Lopez et al., 2005; Lückemann et al., 2016). Importantly, recent findings demonstrated that the administration of low doses of the UCS (10-25 %) shortly after CS re-presentation as *reminder cues*, not only strengthened but also sustained these learned immunosuppressive responses. This strategy has been shown to prolong heart allograft survival and to attenuate disease progression in rat models of brain tumor and rheumatoid arthritis (Hadamitzky et al., 2016; Lückemann et al., 2020; Hetze et al., 2022). To evaluate the potential effectiveness of this dose-reduction strategy in an allergy-related clinical setting, the present study applied a taste-immune associative learning protocol with CsA in a rat model of contact hypersensitivity (CHS).

## Material and Methods

### Animals

Male Dark Agouti rats (DA/HanRj; nine to ten weeks old, 190-220 g; Janvier Labs, France) were group housed in a temperature (20° C) and humidity (55 ± 5 %) controlled facility under a 12/12-h reversed light/dark cycle (lights off at 7:00 a.m.). After acclimatization for two weeks, rats were housed individually with *ad libitum* access to food and tap water. The animal facilities and experimental procedures were in accordance with National Institutes of Health and Association for the Assessment and Accreditation of Laboratory Animal Care guidelines and the ARRIVE guidlines and were approved by the Institutional Animal Care and Use Committee (LANUV TSG-Nr. G1884/21 Düsseldorf, North Rhine-Westphalia).

### Drugs

A stock solution of the immunosuppressive drug CsA was prepared freshly every day by dissolving 200 mg CsA (LC Laboratories, Woburn, USA) in 200 µL 95 % ethanol (Braun, Melsungen, Germany) and 1800 µL Miglyol (Caelo, Hilden, Germany). Corresponding to the animal’s individual weight, this stock solution was further diluted with (0.9 % NaCl, Braun, Melsungen, Germany) to gain the respective drug dosages of 20, 40 and 80 mg/kg body weight (Hadamitzky et al., 2016; Lückemann et al., 2020).

### CHS induction and treatment

CHS was induced using the hapten DNFB (Sigma-Aldrich, St. Louis, USA) in two phases (Manresa, 2021). In a sensitization phase, the hapten (100 µL of 0.5 % DNFB dissolved in 4:1 acetone - olive oil) was applied on the animal’s shaved abdomen on two consecutive days. Following an incubation period of four days, the sensitization phase was initiated, where animals were re-exposed to the allergen on one ear (20 µL of 0.3 % DNFB). The corresponding ear serves as control and was treated with a vehicle solution. 24 h later, ear thickness was measured (**Figure 1 A**). All procedures were performed under anesthesia to prevent stress for the animals (1.5 - 2% isoflurane mixture with oxygen; Isothesia, Piramal Critical Care B.V., Voorschoten, Netherlands).

**Figure 1.**
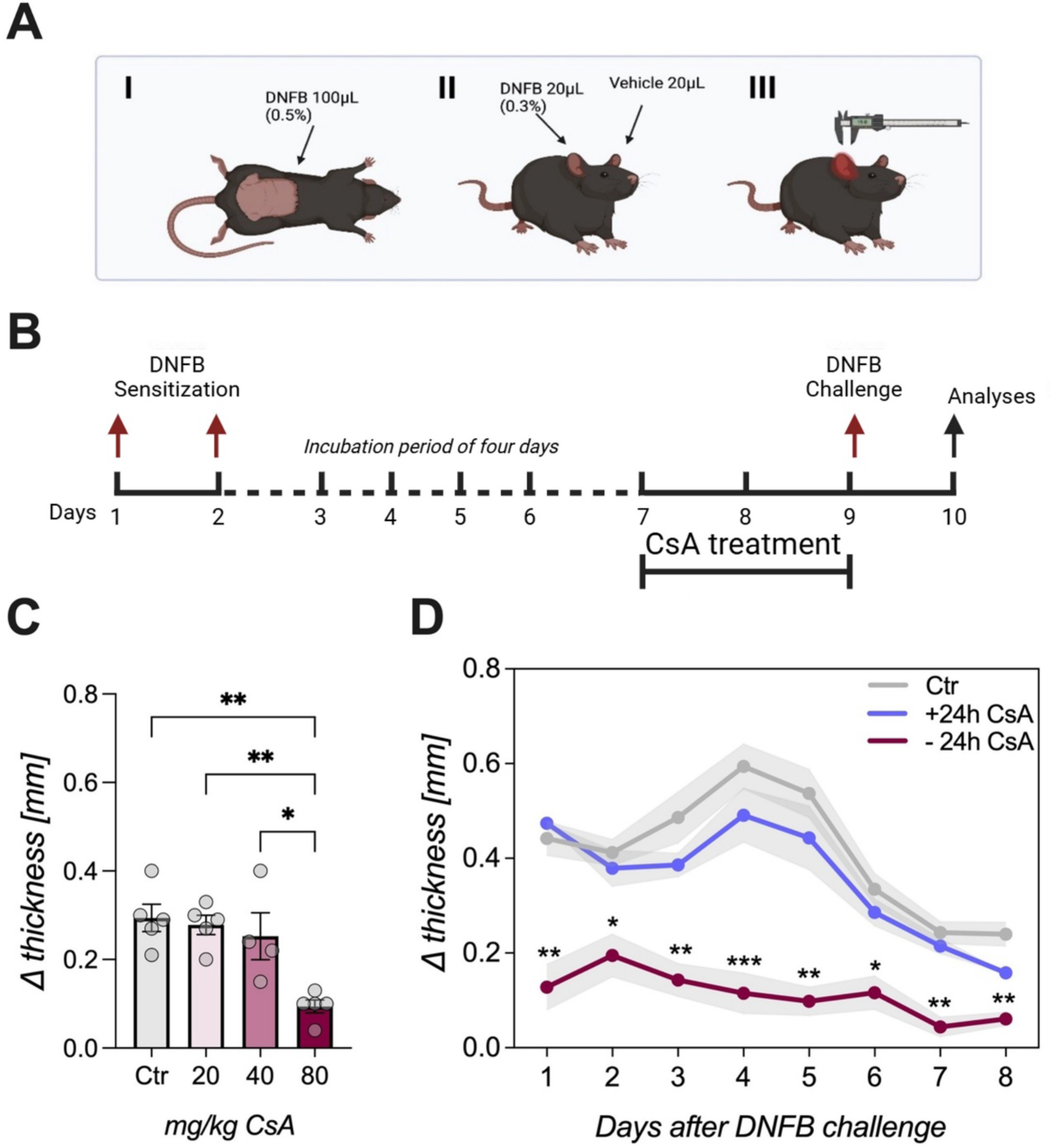
CsA treatment on CHS disease progression. (**A**) Schematic overview of CHS induction. Animals were sensitized on two consecutive days with DNFB (100 µL 0.5 %) on the shaved abdomen. After an incubation time of four days, animals were challenged on one ear with DNFB (20 µL 0.3 %), the other ear serves as control ear. 24 h later ear thickness was measured with a digital caliper. (**B**) Schematic representation of the dose finding design. Animals were sensitized with DNFB on two consecutive days. After an incubation time of four days, animals were treated with different doses of CsA for three consecutive days. On the third day, animals were challenged with DNFB and ear thickness was measured 24 h later. (**C**) Treatment with 80 mg/kg resulted in a significant attenuation of thickness difference (ANOVA followed by Bonferroni post hoc analysis, *p<0.05 **p<0.01; n=4/group). (**D**) To assess the impact of the starting point of CsA (80 mg/kg) treatment, animals were subjected to continuous treatment regimen at different time points, either pre (*-24 h CsA*) or post (*+24 h CsA*) DNFB challenge. Ear swelling was monitored over eight days. Bonferroni post hoc tests revealed significant reduction of ear thickness difference of the *-24 h CsA* group compared to the control group at every time point (*p<0.05, **p<0.01, ***p<0.001; n=5/group). Data are shown as mean ± SEM.

### Ear thickness measurement

Ear thickness of the challenged and the control ear were measured at the middle of the ear lobe using a digital caliper (Kroeplin, Germany). Symptomatology of CHS was analyzed as ear thickness difference between both ears (Δ thickness).

### Dose-response and treatment of CHS

Three different doses of CsA (20, 40 and 80 mg/kg) were applied intraperitoneally (i.p.) and tested regarding their efficacy to prevent CHS symptoms (ear swelling). Animals were sensitized as described above and four days later treatment over three consecutive days was initiated. On the third day, animals were re-exposed to the allergen and ear thickness was measured 24 h later. In addition, treatment effects of CsA on CHS were analyzed starting at two different time points: 24 h before the challenge with the allergen (*-24 h CsA group*) and 24 h after the challenge (*+24 h CsA*). Ear thickness was measured over a period of eight days and compared with a non-treated control group (*Ctr*; **Figure 1 B**).

### Behavioral conditioning protocol - Experiment 1

For the established conditioning paradigm, animals underwent a training phase with access to water in the morning (9 a.m.) and evening (5 p.m.) for 15 min only. After each drinking session, the bottles were weighed to measure fluid intake. Individual mean water consumption of all sessions over a period of five days was taken as baseline level (100 %) for ‘‘normal” fluid intake. After these five training days, acquisition started and animals were randomly allocated into four treatment groups. During three acquisition trials, separated by 72 h, all animals received saccharin and an i.p. injection with CsA (except handling control animals (*Veh*)) in the morning session. In the evening session, all animals received water. Five days after the last acquisition all animals were immunized (induction of CSH as described above) and received water during the morning and evening drinking sessions until day 22 before retrieval was initiated. At retrieval, rats of the conditioned group (*CS)* received saccharin in the morning sessions and water in the evening. To compare conditioning effects with a standard pharmacological treatment, animals of the *US* group received i.p. injections of CsA (80 mg/kg) during the morning session of the retrieval phase. Control animals for residual effects of CsA administration (*CS0*) were neither re-exposed to the saccharin (CS) nor did they receive additional CsA at retrieval. Three retrieval trials were performed with DNFB challenge on the second trial, whereas ear swelling was measured 1 h after the last trial (**Figure 2 A, B**).

**Figure 2.**
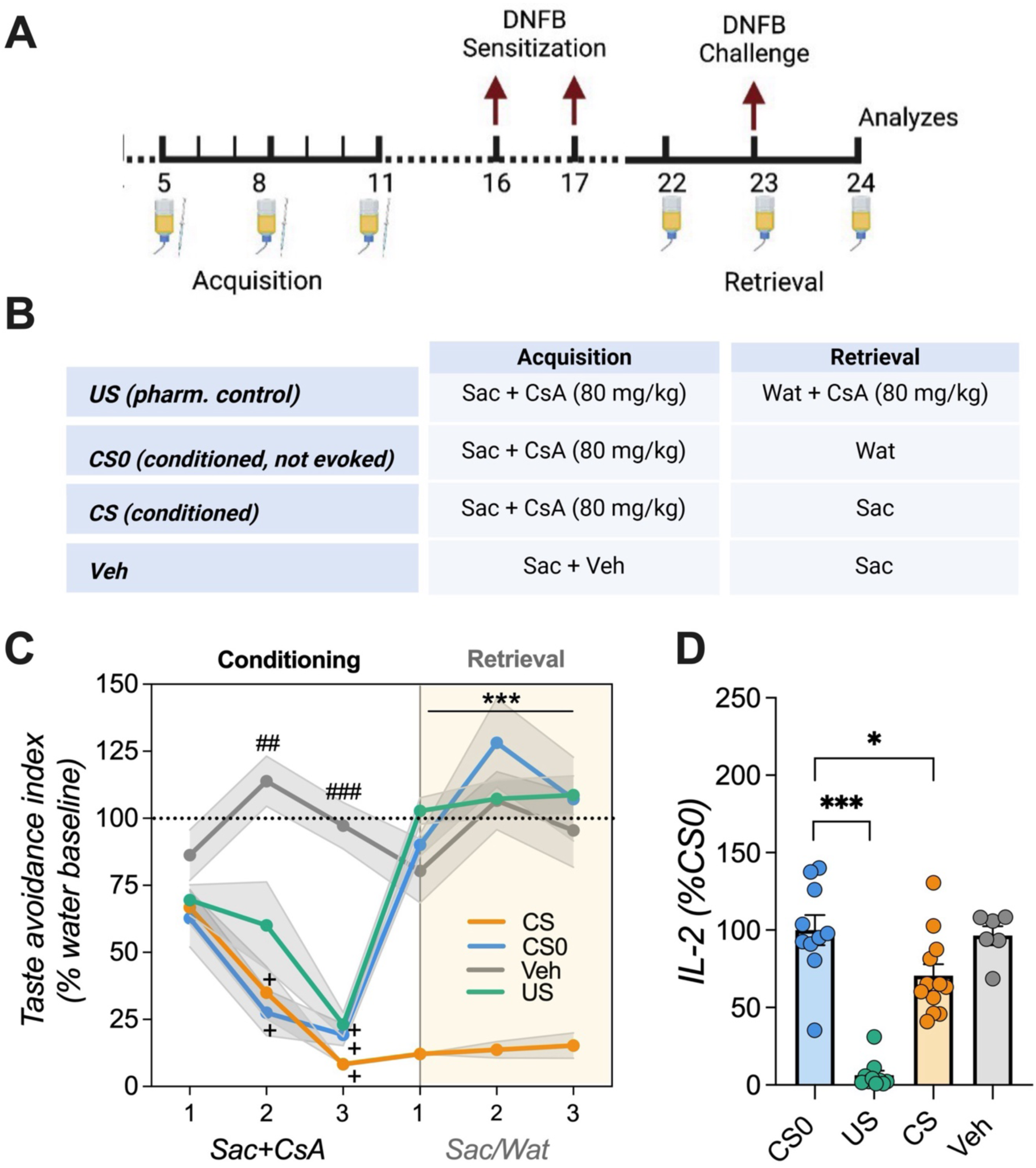
Taste-immune associative learning. (**A**) Schematic representation of conditioning paradigm and (**B**) group allocation. Animals underwent three acquisition trials. During these trials the *CS, US,* and CS*0* groups received saccharin and an injection with CsA (80 mg/kg) prior to sensitization with DNFB. After the incubation time, retrieval started with presentation of the saccharin solution (*CS* and *Veh* group) or water (*US* and *CS0* group). The *US* group additionally received 80 mg/kg CsA. On the second day of retrieval, animals were challenged with DNFB and 24 h later ear thickness was measured. (**C**) Taste avoidance index. Conditioned animals showed a pronounced CTA on the second and third acquisition day and over the course of retrieval, reflected by a significantly lower saccharin consumption (ANOVA followed by Bonferroni post hoc analysis; ##p<0.01; ###p<0.001 = *Veh* vs. all groups; +++p<0.001 = all groups vs. acquisition day 1; ***p<0.001 = *CS* vs. all groups; n=6-12/group). (**D**) IL-2 levels of *ex-vivo* anti-CD3 stimulated splenocytes. Compared to the *CS0* group, *US* and *CS* animals showed reduced IL-2 amounts. Data are presented as percentage of *CS0* group (ANOVA followed by Bonferroni post hoc analysis; *p<0.05, ***p<0.001; n=6-12/group). Data are shown as mean ± SEM.

### Memory-updating of protocol - Experiment 2

The established conditioning paradigm was performed as described above in identically treated *US* and *CS0* groups. However, two additional experimental groups (*CSlow*, *USlow*) were now included in the protocol. Both groups were conditioned with three acquisition trials receiving saccharin as CS and an i.p. injection with CsA (80 mg/kg) as UCS as described in *Experiment 1*. During each of the six retrieval trials, however, the *CSlow* animals received an i.p. injection with sub-effective CsA (20 mg/kg) after saccharin exposure in the morning session. The *USlow* group served for controlling the pharmacological efficacy of the sub-effective doses (“*reminder cues*”) by receiving water at both drinking sessions together with an injection of sub-effective CsA (20 mg/kg) in the morning session. Retrieval was conducted on six consecutive days, with a DNFB challenge on the second day, while ear thickness was measured and samples were collected 1 h after the final retrieval trial (**Figure 4 A, B**).

### Conditioned taste avoidance (CTA)

After each morning and evening drinking session, the total amount of liquid consumed was assessed by weighing the bottles. Saccharin consumption was calculated as a percentage of the individual baseline water consumption.

### Splenocytes isolation

Spleens were collected 1 h after the last retrieval trial and transferred into a falcon with ice cold HBSS (Hank’s Balanced Salt Solution, Gibco®, Life TechnologiesTM, Carlesbad, USA) and disrupted with a syringe plunge. Red blood cells were then lysed with BD Pharm Lyse (BD Pharmingen, Heidelberg, Germany) and splenocytes were washed in cell culture medium (RPMI + 10 % FBS + 0.1 % gentamycin) before being filtered through a 70 μm nylon cell strainer. After washing the suspension, cell concentrations were determined via an automated cell counter (Vet abc, Medical Solution, Steinhausen, Switzerland) and cell numbers were adjusted to a final concentration of 5 x 10^6^ cells/mL. Splenocytes were seeded in a 96 well plate (250.000 cells/well) and stimulated with 1 µg/mL of mouse anti-rat CD3 monoclonal antibody (clone: G4.18, BD Biosciences) for 24 h in a humidified incubator (37°C, 5 % CO2).

### IL-2 ELISA

IL-2 amounts of the supernatants were measured using a sandwich ELISA (Quantikine®ELISA Rat IL2, R&D systems, Minneapolis, USA) according to the manufacturer’s instructions. Optical density was assessed at 540 and 570 nm using Fluostar OPTIMA Microplate Readers (BMG Labtech, Offenbach, Germany). Cytokine concentrations were calculated using a log-log curve-fit standard curve.

### Histology

Histological analysis of the challenged ear was performed to quantify the mast cell infiltration. Ears were collected, cut longitudinally, and fixed in 4 % paraformaldehyde for one week. Tissues were embedded in paraffin and cut serially into slices of 5 µm thickness using a microtome. Sections were stained with 0.01% toluidine blue for one minute and scanned using Scope Scanner (Aperio CS2 Scanner, Leica Biosystems, Deer Park, USA). Three sections per animal with a total of nine areas were counted using *ImageJ* (version: 2.9.0/1.53t; National Institutes of Health) and evaluated in a blinded manner.

### Statistical analysis

Statistical analyses were performed using SigmaPlot (Version 12.3, Systat Software San Jose, CA, USA) and GraphPad (Version 9.5.1, Graph Pad Software, San Diego, CA, USA). Normality was tested via Shapiro-Wilk-Test and data were log transformed when necessary. P value was considered significant at <0.05. Two-way analysis of variance (ANOVA) with *group* (treatment) as one factor and *time* (days) as a within-subjects factor was used to analyze behavioral data as well as ear thickness measurements. When appropriate, post-hoc individual comparisons between groups were determined by Bonferroni t-tests. One-way ANOVA followed by Bonferroni or Dunnett’s Method post hoc analysis was used for all other experiments. The numbers of animals per treatment group, which differ marginally due to technical reasons, are reported in the figure legends.

## Results

### Pre-treatment with high doses of CsA attenuates CHS symptoms

To determine the effective dose of CsA for preventing allergic DNFB-induced ear swelling, a dose response study was conducted, revealing differences between the applied dosages (ANOVA: F(3,15)=9.552; p<0.001). Only animals treated with 80 mg/kg CsA for three consecutive days showed a significant attenuation of ear swelling compared to controls and animals treated with 20 or 40 mg/kg CsA (p<0.05; **Figure 1 C**). Subsequently, different time points for a therapeutic intervention with 80 mg/kg CsA were examined by observing ear swelling over eight days. ANOVA revealed a main effect for the factor *time* (F(7,112=54.24; p<0.001), *group* (F(3,112)=23.02; p<0.001) and a *time x group* interaction (F(21,112)=4.734; p<0.001). Noteably, only pre-treatment that started 24 h before challenge led to a significantly diminished ear thickness compared to untreated controls and the group receiving CsA treatment 24 h post challenge (*Ctr* vs *-24 h CsA* p<0.05). In control animals, ear thickness increased peaking on day four (**Figure 1 D**).

### Taste-immune associative learning in CHS

In a first setup (*Experiment 1*), animals underwent three conditioning sessions where a saccharin solution (CS) was paired with CsA injections (UCS; 80 mg/kg), before being immunized with DNFB on two consecutive days. Since pre-treatment (24 h before immunization) was necessary to prevent ear swelling, retrieval of conditioned immunosuppression began one day prior to DNFB challenge (**Figure 2 A, B**). Conditioned animals showed a pronounced CTA across all three retrieval trials (ANOVA: for *group* (F(3,68)=80.486; p<0.001) and *time* (F(2,68)=6.375; p<0.01), reflected by significantly diminished fluid intake in the *CS* group (**Figure 2 C**; p<0.001). Concurrently, IL-2 cytokine production of *ex-vivo* anti-CD3 stimulated splenocytes differed significantly among groups (ANOVA F(3,34)=33.54; p<0.001; **Figure 2 D**). Post-hoc analysis revealed that IL-2 levels in conditioned *(CS)* and fully treated (*US)* animals were significantly lower than in the *CS0* group (p<0.05, p<0.001 ANOVA revealed significant groups differences of allergic symptoms (ANOVA: F(3,34)=44.46; p<0.001), but only animals treated with 80 mg/kg CsA exhibited reduced ear swelling (p<0.001; **Figure 3 A**). Consistent with IL-2 production, histological staining revealed increased mast cell infiltration at the inflammation site in US and CS groups compared to the controls (**Figure 3 B**). However, correlation analysis demonstrated a significant negative interaction between ear thickness and mast cell density (R squared=0.127, p=0.028; **Figure 3 C, D**). Analysis of immune cell subsets such as B cells, MHC-II+ cells, and APCs in cervical and axillary lymph revealed reductions only in the *US* but not conditioned group (Supplementary Figure 1).

**Figure 3.**
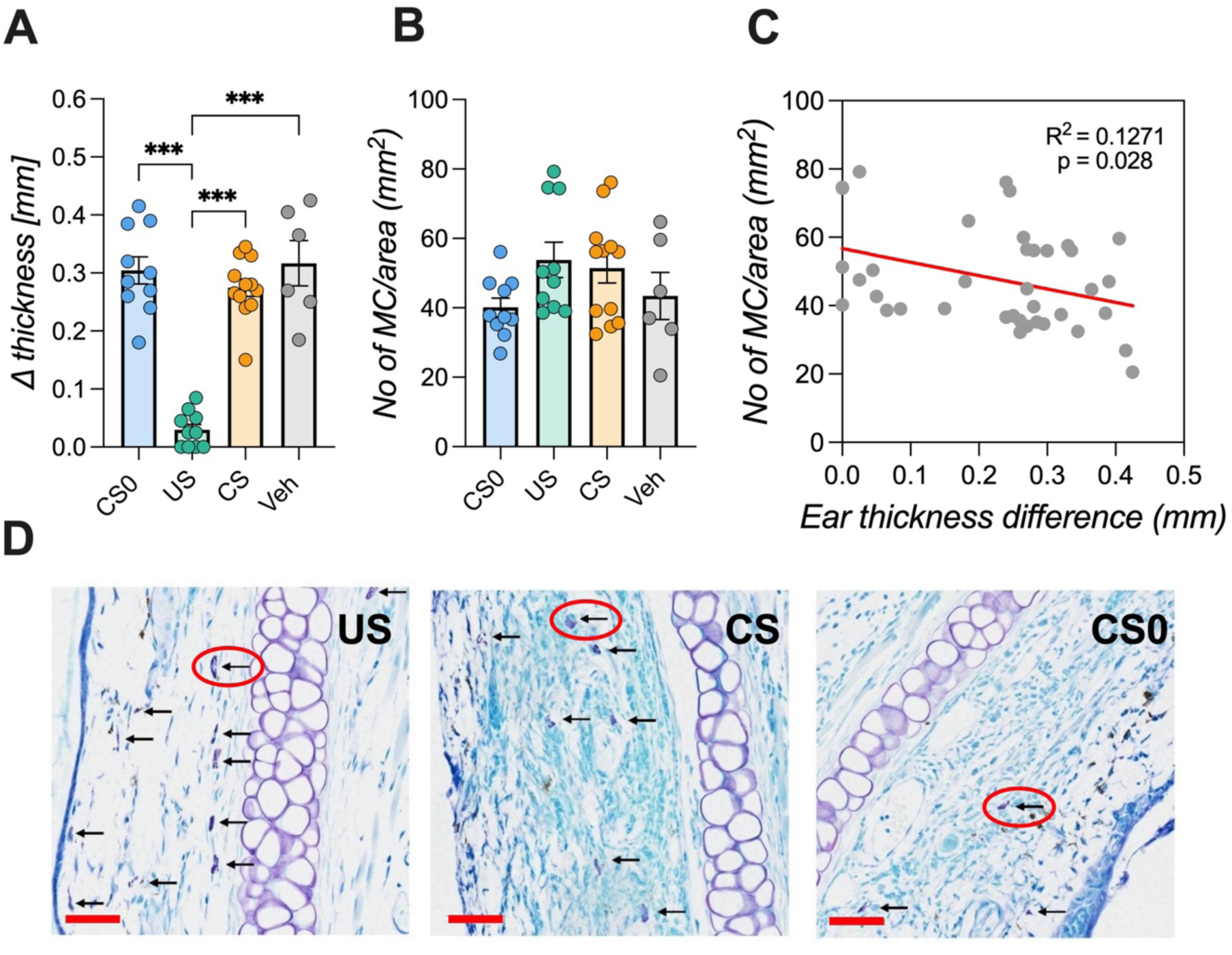
Local CHS inflammation following taste-immune associative learning. (**A**) Ear thickness was measured 1 h after the last retrieval trial. Animals receiving the full pharmacological dose of CsA (*US* group) showed significantly attenuated ear swelling compared to the other groups (ANOVA; Bonferroni post hoc analysis, ***p<0.001; n=6- 12/group). (**B**) Mast cell infiltration in the ear was counted and standardized to the area. *US* and *CS* groups showed a tendency towards higher numbers of mast cells per area but did not reach level of significance. (**C**) Correlation analysis between ear thickness difference and mast cell infiltration showed a significant negative interaction (Pearsons’s correlation, R squared=0.127; p=0.028). Data are shown as mean ± SEM. (**D**) Representative images of a toluidine-stained ears (5 µm section, 50 µm red scale bar). Mast cells (arrows) and cartilage are visible in violet. Sections of *US* and *CS* animals demonstrate a higher amount of mast cell infiltration into the site of local allergic reaction compared to the *CS0* group.

### Memory-updating of taste-immune associative learning boosts systemic immunosuppression but does not affect local CHS symptoms

In the second setup (*Experiment 2*), animals underwent three conditioning trials using CsA (80 mg/kg) as UCS and saccharin as CS before being immunized with DNFB. Similar to *Experiment 1*, retrieval began one day prior to DNFB challenge (**Figure 4 A**). However, conditioned animals in the *CSlow* group received saccharin paired with injections of 25 % (20 mg/kg) of the full therapeutic CsA dose. Control groups received water together with injections of either 20 mg/kg CsA (*USlow* group) or 80 mg/kg CsA (*US* group), or vehicle (*CS0,* **Figure 4 B**). ANOVA revealed significant *group* (F(3,90)=152.634; p<0.001) and *time* effects (F(5,90=3.78; p<0.01) at retrieval. The *CSlow* group exhibited a robust CTA over all six retrieval trials as reflected by significantly reduced fluid intake (p<0.001; **Figure 4 C**). Systemic IL-2 cytokine levels were also affected (F(3,18)=12,85; p<0.001) with significantly reduced IL- 2 levels in the CSlow and US (80 mg/kg CsA) groups compared to CS0 animals (p<0.05, p<0.001; **Figure 4 D**). In line with *Experiment 1*, reduced ear swelling reduction was observed only in the US group but not in conditioned animals (ANOVA: F(3,18)=4.02; p<0.05; **Figure 5 A**). Furthermore, local immune cell infiltration did not differ significantly between groups (**Figure 5 B**).

**Figure 4.**
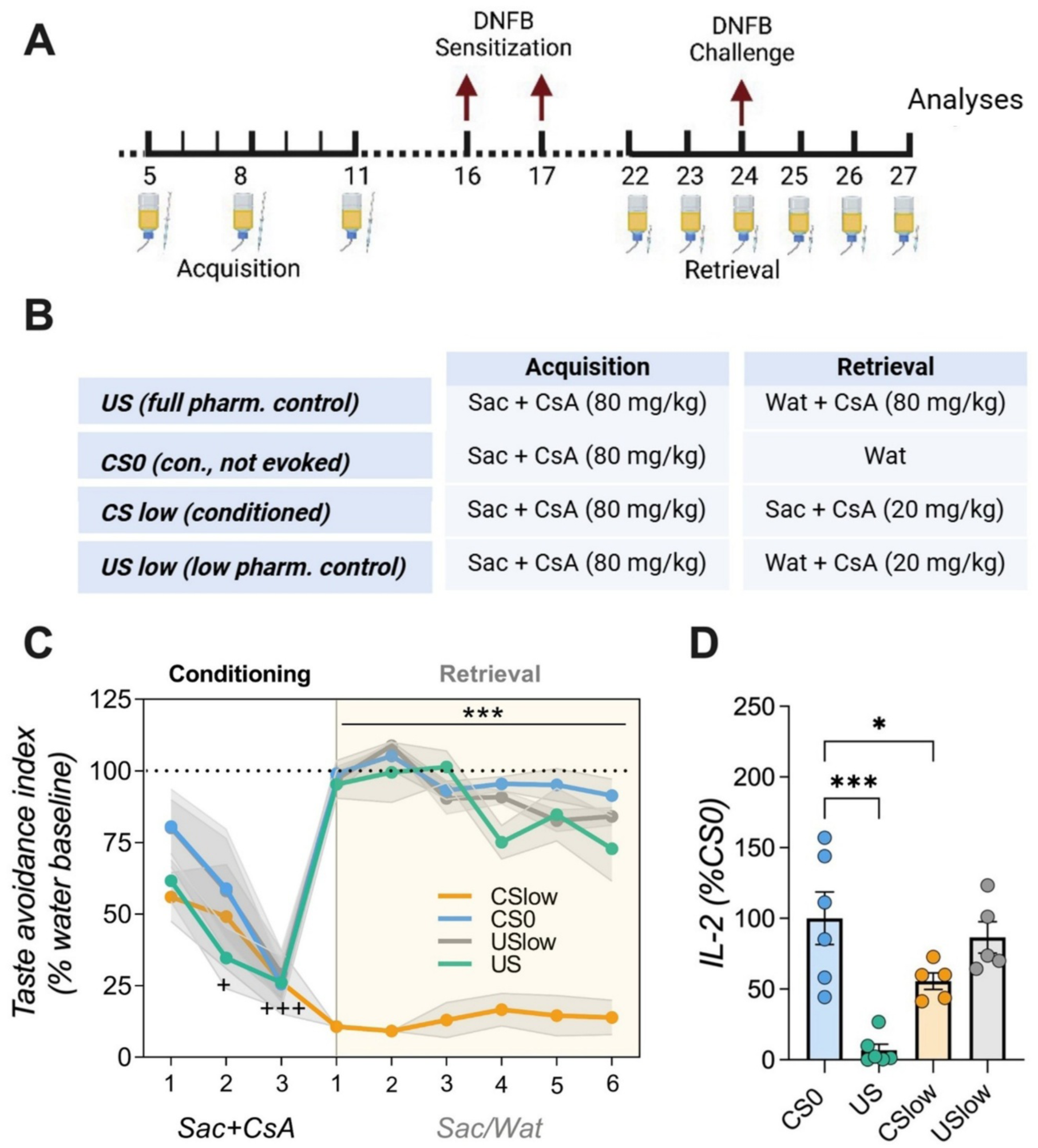
Taste-immune associative learning with reminder cues. (**A**) Schematic representation of conditioning paradigm and (**B**) group allocation. During three acquisition trials (CS/UCS parings), animals of all groups (*CSlow*, *USlow*, *US*, *CS0*) received saccharin and an injection with CsA (80 mg/kg). Subsequently, animals were sensitized with DNFB. 5 days following sensitization, conditioned animals (*CSlow*) received saccharin paired with injections of 25 % (20 mg/kg) of the full therapeutic CsA dose during retrieval. Control groups received either water together with injections of either 20 mg/kg CsA (*USlow* group) or 80 mg/kg CsA (*US* group), or vehicle (*CS0* group). Animals were challenged with DNFB on the second day of retrieval. (**C**) Taste avoidance index. Compared to all other groups, conditioned animals (*CSlow*) displayed a pronounced CTA, over the course of retrieval (ANOVA followed by Bonferroni post hoc analysis; +p<0.05, +++p<0.001 = all groups vs. acquisition day 1; ***p<0.001 = *CSlow* vs. all groups; n=5-6/group). (**D**) *US* and *CSlow* groups also significantly differed in IL-2 production of *ex-vivo* anti-CD3 stimulated splenocytes compared to *CS0.* Data are presented as percentage of *CS0* group. (ANOVA followed by Dunnett’s post hoc analysis, +p<0.05; ++p<0.001; *p<0.05, ***p<0.001; n=5-6/group; Data are shown as mean ± SEM).

**Figure 5.**
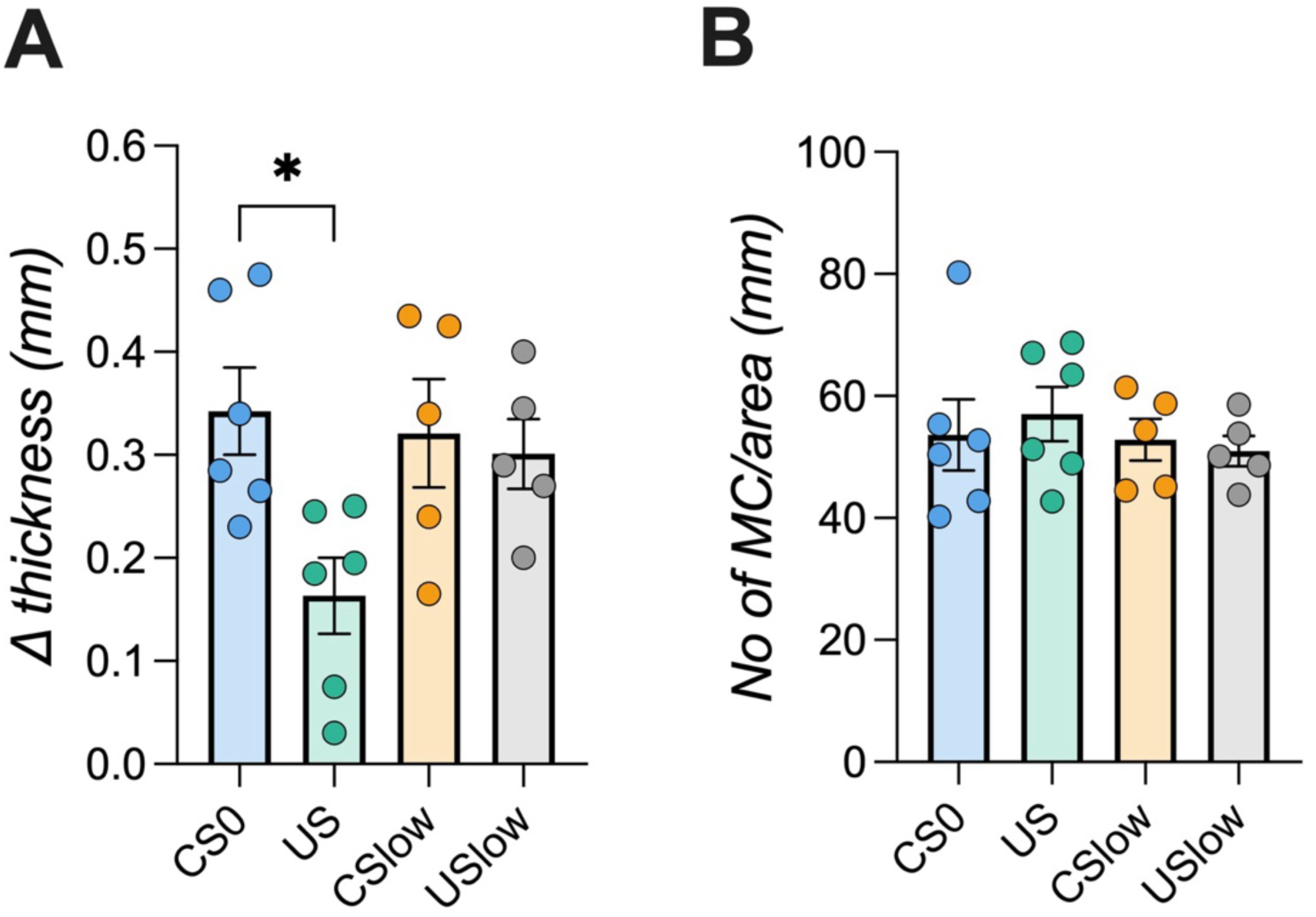
Local CHS inflammation following taste-immune associative learning with reminder cues. (**A**) Ear thickness measurement revealed significant diminished swelling in *US* animals compared to the *CS0* group (ANOVA followed by Bonferroni post hoc analysis, *p<0.05). (**B**) Mast cell infiltration into the inflamed ear was analyzed and no differences were observed. Data are shown as mean ± SEM (n=5-6/group).

## Discussion

The present study investigated the potential of taste-immune associative learning with CsA in a rat model of CHS. Compared to controls, we demonstrated that conditioned immunopharmacological responses with CsA, reflected by suppressed level of the proinflammatory cytokine IL-2, were preserved when 25 % of the drug dose (20 mg/kg CsA) was applied at retrieval alongside with the CS (saccharin). However, this conditioned immunosuppression was insufficient to prevent the emergence of local allergic ear swelling.

The chronic use of small-molecule immunosuppressants can lead to significant side effects (Bosche et al., 2015), highlighting the urgent need for alternative or supportive treatments options in immunological diseases. Based on the intense crosstalk between the central nervous system and the peripheral immune system (Pavlov and Tracey, 2017; Koren et al., 2021), conditioning of immunopharmacological effects via associative learning has been suggested as means to reduce drug dosages and associated side effects, while maintaining treatment efficacy (Hadamitzky and Schedlowski, 2022). Numerous experiments in rodents document the potential clinical relevance of taste-immune associative learning protocols by reporting symptom alleviation in various animal disease models (Hadamitzky et al., 2020). Against this background, we applied an established paradigm of conditioned immunosuppression with CsA in a DNFB-induced CHS rat model. One major difference from previous studies was, however, the necessity of using a high dose of CsA (80 mg/kg) to prevent DNFB-induced ear swelling, compared to the standard 20 mg/kg. This requirement aligns with other studies reporting the need for high doses in similar contexts (Cramer et al., 1992; Natsuaki et al., 2002; Manresa et al., 2019; Fukushima et al., 2020; Li et al., 2020). Consequently, the taste-immune associative learning paradigm was modified to incorporate high dose CsA as UCS. While earlier studies exclusively employing 20 mg/kg CsA for taste- immune associative learning, our findings showed that conditioned animals exhibited pronounced CTA and reduced IL-2 cytokine levels in ex vivo CD3-stimulated splenocytes. Nonetheless, analysis of the local reaction site revealed no significant impact of immune conditioning. Despite a trend towards higher mast cell infiltration, only the pharmacological group treated with 80 mg/kg CsA exhibited reduced ear thickness.

CsA is known to inhibit mast cells’ histamine release (Sperr et al., 1996; Toyota et al., 1996; Kovalik et al., 2012), potentially accounting for the higher cell count observed in these groups, whereas the reduced numbers in controls might reflect a degranulated stage. Given that histamine release contributes to vasodilation-dependent oedema (Ashina et al., 2015; Elieh Ali Komi et al., 2020), its inhibition in CsA treated animals likely resulted in diminished ear thickness. However, in conditioned animals the trend towards higher of mast cell numbers as insufficient to reduce swelling, likely due to the complex immune responses involved in CHS, which comprise various cytokines and immune cells (Scheinman et al., 2021). Analyses of these mediators showed reductions in CsA treated but not conditioned animals (data not shown). Interestingly, a similar study using a dinitrochlorobenzene (DNCB)-induced CHS model reported a conditioned reduction in ear swelling with only 20 mg/kg CsA (Exton et al., 2000). This discrepancy underscores the allergen-dependent variability in allergic responses (Wu et al., 2022). Thus, DNFB-induced reactions may have been too severe to respond to conditioned immunosuppressive effects at the local site of allergic reaction.

Conditioned responses are known to weaken over time with repeated exposure to the CS without the UCS (Berman and Dudai, 2001). To address this, previous studies introduced a non-invasive memory-updating approach where low or sub-effective doses of the drug used as UCS are paired with the CS during retrieval, preventing extinction of the learned immune memory. (Hadamitzky et al., 2016; Lückemann et al., 2021). This approach has demonstrated extended graft survival in transplantation (Hadamitzky et al., 2016), reduced rheumatoid arthritis symptoms (Lückemann et al., 2020) and attenuated glioblastoma disease progression (Hetze et al., 2022) in animals. In the present study, 25 % of the initial drug dose was administered together with the CS during retrieval. Conditioned animals exhibited a pronounced CTA across six retrieval trials. Compared to the previous experiment, splenic proinflammatory IL-2 cytokine levels were further reduced, indicating a strengthened CS-UCS association. Despite this enhancement, CHS symptomatology including allergic ear swelling and histological findings, remained unaffected. The lack of histological differences might reflect the timing of analysis, as the allergic reaction may have entered a resolution phase by day four post-challenge (Scheinman et al., 2021). The unaltered symptoms could be attributed to the intensity of the DNFB-induced allergic response, which may have been too strong for conditioned immunosuppressive effects to overcome. Similar findings were observed in a murine model of autoimmune uveitis, where conditioned reductions in IL-2 cytokine levels did not translate to symptom improvement (Bauer et al., 2017). It was suggested that active sensitization might mask conditioned symptom reduction, and splenocyte transfer from conditioned to naïve animals could help reveal these effects. Such approaches may also clarify the results of the current study. It is important to note that CsA is not a first-line treatment for CHS, nor is pre-treatment a standard clinical approach. Thus, future studies should investigate the use of alternative drugs, such as dexamethasone, and explore adjusted treatment schedules. Additionally, subjective symptoms like itch, which better reflect patient burden, should be prioritized over tissue swelling. Previous studies have shown reduced itch despite unaffected wheal sizes (Darragh et al., 2015).

Taken together, the present study confirms that conditioned immunosuppression can be preserved by using a memory-updating approach. However, despite a strengthened CS- UCS association CHS symptomatology and histology were unaffected in conditioned animals. These results highlight the need to further investigate the underlying mechanisms and clinical applicability of taste-immune associative learning approaches to optimize its responsiveness towards disease-specific symptoms and to enhance its translational potential.

## Acknowledgements

We thank Falk Kaehler, Fabian Tegelhütter and Jasmin Schmidt for their technical assistance.

## Author contribution

Yasmin Salem, Stephan Leisengang, Marie Jakobs, Kirsten Dombrowki, Julia Bihorac, Laura Heiss-Lückemann Sebastian Wenzlaff, Lisa Trautmann, and Martin Hadamitzky conducted the experiments. Yasmin Salem, Manfred Schedlowski, Tim Hagenacker, and Martin Hadamitzky. designed the study. and Yasmin Salem and Martin Hadamitzky wrote the manuscript. Yasmin Salem performed all analyses. Martin Hadamitzky and Manfred Schedlowski acquired the financial support for the project leading to this publication. All authors discussed and revised the manuscript.

## Data Availability declaration

All data supporting the findings of this study are available within the paper and its Supplementary Information.

