## Supplemental Figure 1 for "Sustained learned immunosuppression could not prevent local allergic ear swelling in a rat model of contact hypersensitivity"

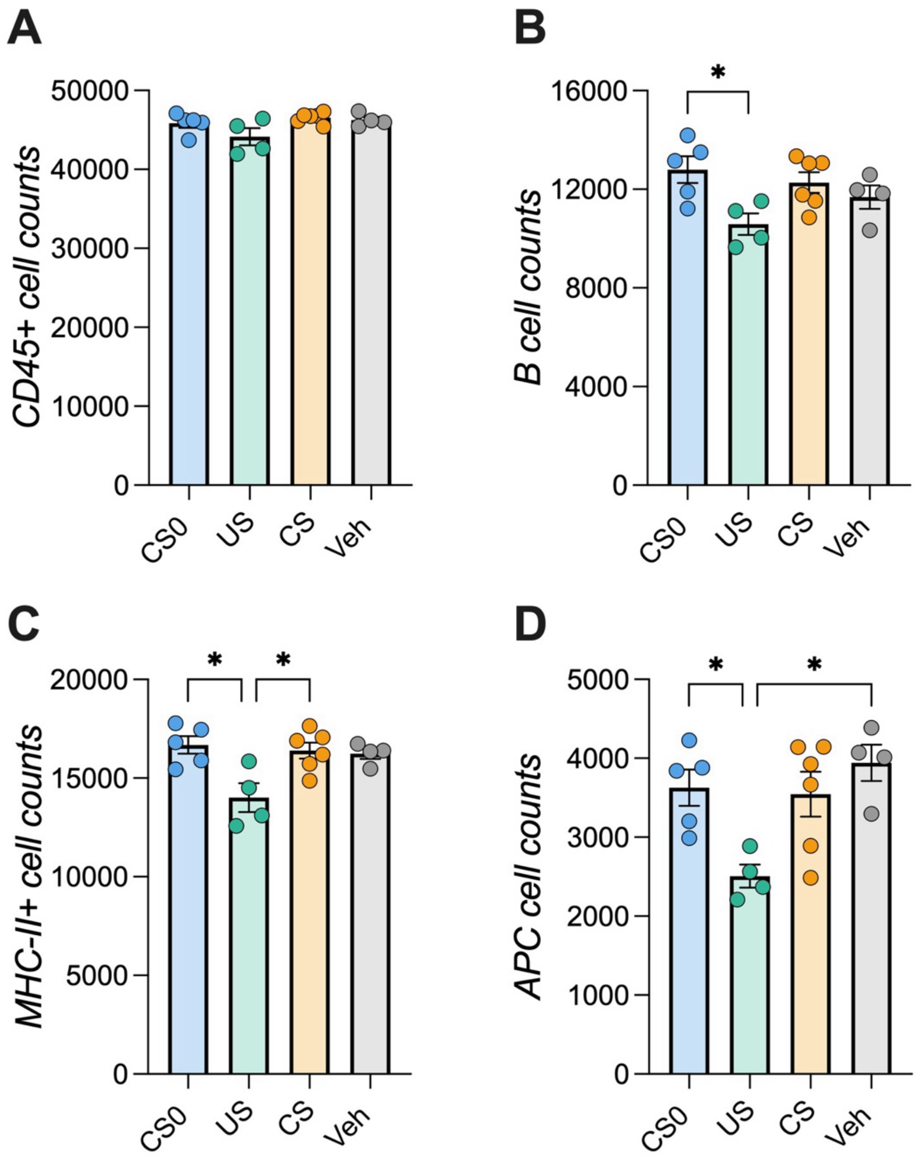


**Supplementary Figure 1. Immune cell subset distribution of draining lymph nodes.** Immune cells of draining lymph nodes (axillary and cervical) were analyzed via flow cytometry. No differences between groups were observed in (**A**) CD45+ cells. A reduction of numbers in the *US* group for (**B**) B-cells (vs *CS0*), (**C**) MHC-II+ (vs *CS0* and *CS*) and (**D**) APCs (vs *CS0* and *Veh*) were shown (ANOVA followed by Bonferroni post hoc analysis, *$p<0.05$; n=4-6/group). Data are shown as mean ±SEM.
